# Testing the causal impact of amyloidosis on total Tau using a genetically informative sample of adult male twins

**DOI:** 10.1101/2024.07.23.602498

**Authors:** Nathan A. Gillespie, Michael C. Neale, Matthew S. Panizzon, Ruth E. McKenzie, Xin M. Tu, Hong Xian, Chandra A. Reynolds, Michael J. Lyons, Robert A. Rissman, Jeremy A. Elman, Carol Franz, William S. Kremen

**Author notes:** Shared senior authorship. **Corresponding Author** Nathan A. Gillespie, PhD, Virginia Institute for Psychiatric and Behavioral Genetics of VCU, Box 980126, Richmond, VA 23298-0126. **Statement of work** The manuscript is original research. It not been previously published and has not been submitted for publication elsewhere while under consideration.

## Abstract

**INTRODUCTION:** The amyloid cascade hypothesis predicts that amyloid-beta (Aβ) aggregation drives tau tangle accumulation. We tested competing causal and non-causal hypotheses regarding the direction of causation between Aβ40 and Aβ42 and total Tau (t-Tau) plasma biomarkers.

**METHODS:** Plasma Aβ40, Aβ42, t-Tau, and neurofilament light chain (NFL) were measured in 1,035 men (mean = 67.0 years) using Simoa immunoassays. Genetically informative twin modeling tested the direction of causation between Aβs and t-Tau.

**RESULTS:** No clear evidence that Aβ40 or Aβ42 directly causes changes in t-Tau was observed; the alternative causal hypotheses also fit the data well. In contrast, exploratory analyses suggested a causal impact of the Aβ biomarkers on NFL. Separately, reciprocal causation was observed between t-Tau and NFL.

**DISCUSSION:** Plasma Aβ40 or Aβ42 do not appear to have a direct causal impact on t-Tau. In contrast, Aβ aggregation may causally impact NFL in cognitively unimpaired men in their late 60s.

## Introduction

According to the amyloid cascade hypothesis [1], amyloid-beta (Aβ) aggregation drives accumulation of tau tangles, resulting in synaptic dysfunction, neurodegeneration and progression to cognitive decline. This implies a causal link from Aβ aggregation to tau tangles. To our knowledge this link has not been empirically tested using genetically informative direction-of-causation modeling.

We previously explored the heritability of blood-based biomarkers related to risk of Alzheimer’s Disease in a population-based sample of early old-age men [2]. Additive genetic influences explained 44% to 52% of the total variances in Aβ42, Aβ40, total tau (t-Tau), and neurofilament light chain (NFL), a marker of neurodegeneration. All remaining variances were explained by non-shared environmental influences. Since Aβ aggregation was best explained by genetic and non-shared environmental influences [2], if either Aβ phenotypically causes t-Tau, then significant genetic *and* environment covariance should be observed between Aβ and t-Tau biomarkers. Instead, we found that both Aβ42 and Aβ40 were genetically uncorrelated with t-Tau. While this is consistent with there being no causal association between Aβ and t-Tau, we apply a validated means [3] of empirically testing competing hypotheses regarding the direction of causation.

### Aim

Without randomized control trials, Mendelian Randomization or longitudinal data, testing causality between complex traits is difficult. However, by analyzing genetically informative twin data and leveraging the expected differences in the patterns of cross-twin cross-trait correlations, it is possible to falsify hypotheses about the direction of causation between two variables measured on a single occasion [3-7]. Using this approach, we tested competing hypotheses regarding the direction of causation between Aβ and t-Tau plasma biomarkers. We also included exploratory analyses modelling the direction of causation between Aβ and NFL.

## Methods

### Subjects

For detailed sample and data description see Gillespie et al. [2]. The present study comprised of men from the Vietnam Era Twin Study of Aging (VETSA) who participated in a third assessment wave (mean age=68.2, SD=2.5, range=61.4 to 73.3) when plasma biomarkers were examined [2].

### Blood-based biomarker data

Blood was collected under fasting conditions before acquisition and storage at -80°C. The Simoa Human Neurology 3-plex A (N3PA) Immunoassay was used to measure Aβ40, Aβ42, and t-Tau, while the Simoa NF-light assay was used to measure NFL (Quanterix™, Billerica, MA, USA). Biomarkers were regressed onto age at assessment, testing site, storage time, self-reported race/ethnicity, and whether or not twins pairs were assessed on the same day. Residual scores were calculated using the umx_residualize function [8]. Next, the data were normal ranked in R_4.0.3_ [9] and absolute values greater than three standard deviations (SDs) were eliminated to reduce skew. This eliminated a total of 12, 8, 12 and 15 subjects with Aβ40, Aβ42, NFL and t-Tau data respectively. Depending on the biomarker, there were between 988 to 1035 individuals (58% monozygotic and 42% dizygotic twins) with complete data.

### Statistical analyses

Based on biometrical genetic methods [10], the OpenMx_2.20.6_ software package [11] with the raw data Full Information Maximum Likelihood (FIML) option and NPSOL optimizer in R_4.2.2_ [9] was used to decompose the total variance in each biomarker into additive genetic (A), shared or common (C) environment, and non-shared or unique (E) environmental influences while testing competing causal and non-causal hypotheses [10] (see Supplementary).

Illustrated in Figure 1, our null hypothesis predicted that associations between the Aβ and t-Tau biomarkers were explained by correlated, non-causal genetic and environmental influences (for brevity, only genetic influences are shown). We analyzed Aβ40 and Aβ42 separately and tested four competing, nested hypotheses: (**b**) Aβ causes t-Tau via β_1_; (**c**) t-Tau causes Aβ via β_2_; (**d**) reciprocal causation between Aβ and t-Tau via β_1_ and β_2_; and (**e**) no association i.e. β_1 =_ β_2_ = 0. We also modelled the joint impact of both Aβs on t-Tau (see Supplementary Figure S1), followed by exploratory causal modelling between the Aβs and NFL, and finally between t-Tau and NFL.

**Figure 1.**
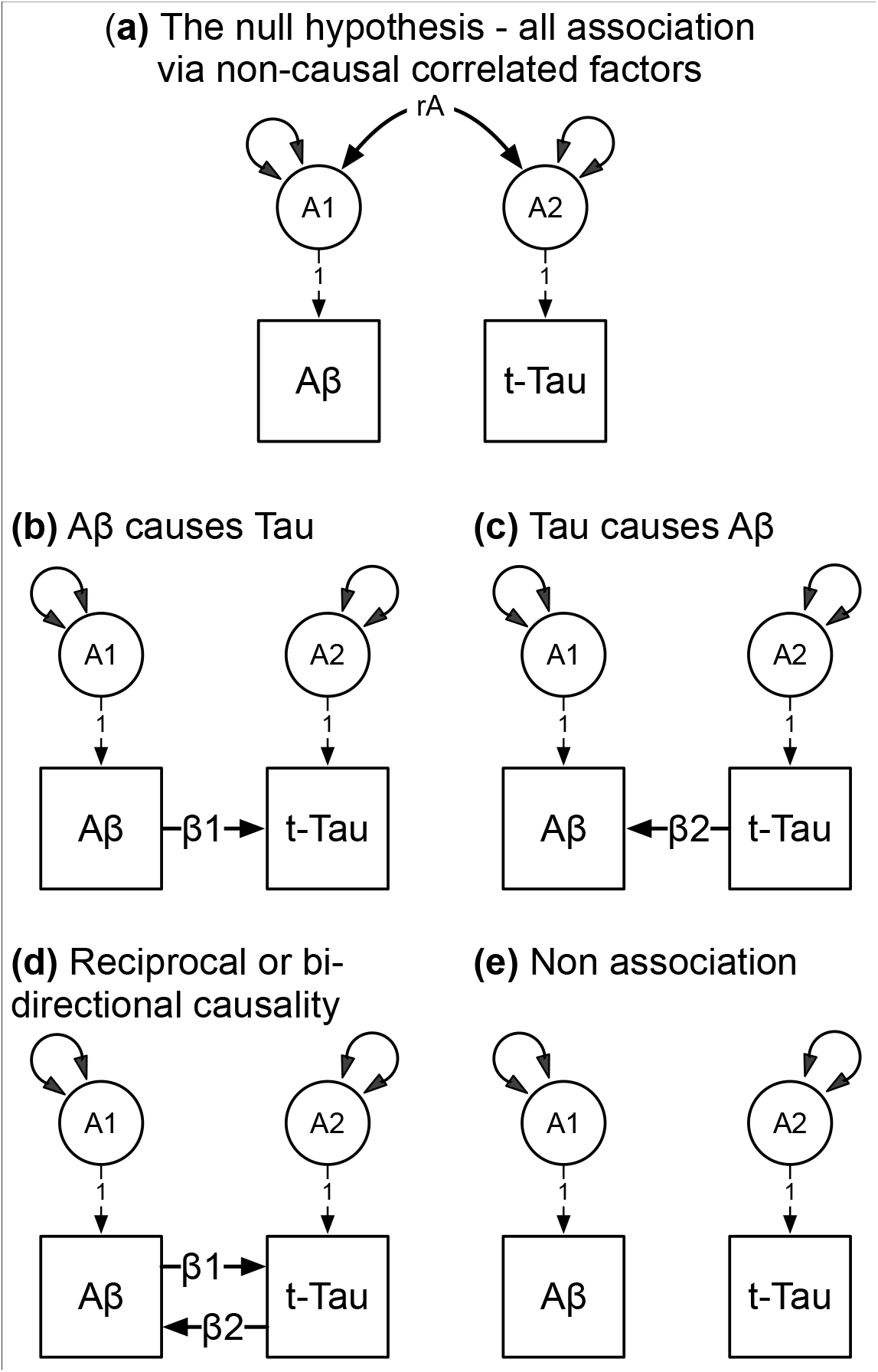
Competing hypothetical models to account for the association between the Aβ and t-Tau biomarkers. **Footnote:** A1 and A2 are latent additive genetic influences for the Aβ and t-Tau plasma biomarkers respectively. Double-headed arrows denote the A1 and A2 genetic variances, and non-causal additive covariance (rA). Latent environmental influences, including residuals and means not illustrated for brevity. Competing models: **(a)** non-causal association stemming from correlated additive genetic influences (rA); **(b)** uni-directional Aβ causes t-Tau via the regression coefficient β1; **(c)** uni-directional t-Tau causes Aβ via the regression coefficient β2; (**d**) reciprocal or bi-directional causation between Aβ and t-Tau; and **(e)** no association between Aβ and t-Tau where β1 = β2 = 0.

The goodness of fit for each model was determined using the likelihood ratio statistic, which is the change in the minus two log-likelihood between the null and each competing model. This statistic, Δ-2LL, is asymptotically distributed as chi-squared with degrees of freedom equal to the difference in the number of free parameters between the null and each competing model. Our determination of the best-fitting model was also based on the optimal balance of complexity and explanatory power using Akaike’s Information Criterion (AIC) [12].

## Results

Model fit comparisons are shown in Table 1. For each set of analyses (i, ii & iii) the null hypothesis predicted that any observed association between Aβ and Tau was attributable to non-causal, correlated genetic and environmental factors.

**Table 1.**
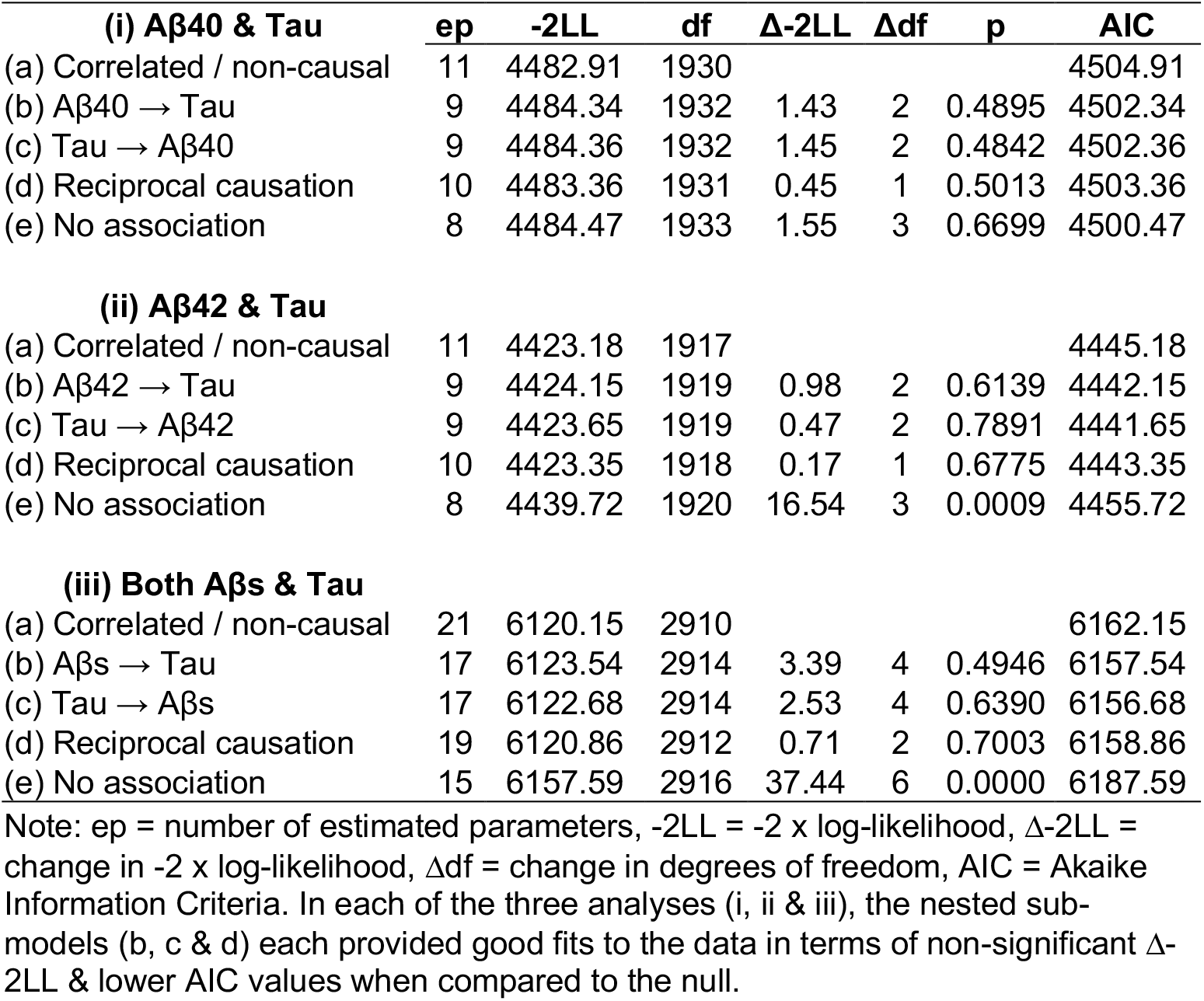
Multivariate model fitting comparisons between the non-causal correlated factors reference model (a) and the two causal (b-c), bi-directional or reciprocal causation (d), and the no association (e) models.

| **(i) Aβ40 & Tau** | **ep** | **-2LL** | **df** | **Δ-2LL** | **Δdf** | **p** | **AIC** |
| --- | --- | --- | --- | --- | --- | --- | --- |
| (a) Correlated / non-causal | 11 | 4482.91 | 1930 |  |  |  | 4504.91 |
| (b) Aβ40 → Tau | 9 | 4484.34 | 1932 | 1.43 | 2 | 0.4895 | 4502.34 |
| (c) Tau → Aβ40 | 9 | 4484.36 | 1932 | 1.45 | 2 | 0.4842 | 4502.36 |
| (d) Reciprocal causation | 10 | 4483.36 | 1931 | 0.45 | 1 | 0.5013 | 4503.36 |
| (e) No association | 8 | 4484.47 | 1933 | 1.55 | 3 | 0.6699 | 4500.47 |
| **(ii) Aβ42 & Tau** |  |  |  |  |  |  |  |
| (a) Correlated / non-causal | 11 | 4423.18 | 1917 |  |  |  | 4445.18 |
| (b) Aβ42 → Tau | 9 | 4424.15 | 1919 | 0.98 | 2 | 0.6139 | 4442.15 |
| (c) Tau → Aβ42 | 9 | 4423.65 | 1919 | 0.47 | 2 | 0.7891 | 4441.65 |
| (d) Reciprocal causation | 10 | 4423.35 | 1918 | 0.17 | 1 | 0.6775 | 4443.35 |
| (e) No association | 8 | 4439.72 | 1920 | 16.54 | 3 | 0.0009 | 4455.72 |
| **(iii) Both Aβs & Tau** |  |  |  |  |  |  |  |
| (a) Correlated / non-causal | 21 | 6120.15 | 2910 |  |  |  | 6162.15 |
| (b) Aβs → Tau | 17 | 6123.54 | 2914 | 3.39 | 4 | 0.4946 | 6157.54 |
| (c) Tau → Aβs | 17 | 6122.68 | 2914 | 2.53 | 4 | 0.6390 | 6156.68 |
| (d) Reciprocal causation | 19 | 6120.86 | 2912 | 0.71 | 2 | 0.7003 | 6158.86 |
| (e) No association | 15 | 6157.59 | 2916 | 37.44 | 6 | 0.0000 | 6187.59 |

Note: ep = number of estimated parameters, -2LL = -2 x log-likelihood, Δ-2LL = change in -2 x log-likelihood, $\Delta$ df = change in degrees of freedom, AIC = Akaike Information Criteria. In each of the three analyses (i, ii & iii), the nested sub-models (b, c & d) each provided good fits to the data in terms of non-significant Δ-2LL & lower AIC values when compared to the null.

When testing the ‘Aβ40 causes Tau’ hypothesis, all four competing hypotheses (both uni-directional, the reciprocal, and the no association model) provided a good fit to the data in terms of non-significant changes in chi-square, whereas the ‘no association’ hypothesis provided the lowest AIC.

When testing the ‘Aβ42 causes Tau’ hypothesis, the changes in chi-square for all three competing causal hypotheses were non-significant, whereas the AIC was lowest for the ‘Tau causes Aβ42’ hypothesis. Note that the ‘no association’ hypothesis could be rejected based on the significant change in chi-square and higher AIC value relative to the three other competing hypotheses.

When testing the ‘Aβ40 & Aβ42 (combined) cause Tau’ hypothesis, the changes in chi-square for each of the three causal hypotheses were again non-significant. However, very little separated their corresponding AIC values. Note that the ‘no association’ hypothesis could again be rejected in terms of the significant chi-square change and highest AIC.

In exploratory analyses, we modelled the multivariate impact of Aβ40 and Aβ42 (Aβs) on NFL (Supplementary Table S1). Among the competing hypotheses, the uni-directional ‘Aβs cause NFL’ hypothesis provided a marginally better fit to the data as judged by the non-significant change in chi-square and lowest AIC. Finally, when modelling the association between t-Tau and NFL, the reciprocal causation model provided a (marginally) best fit to the data, followed next by the ‘t-Tau causes NFL’ hypothesis.

## Discussion

To our knowledge, this is the first genetically informative test of the direction of causation between blood-based biomarkers related to Alzheimer’s Disease. We found no unequivocal support for a causal impact of either Aβ40 or Aβ42 on t-Tau. Instead, alternative uni-directional and reciprocal hypotheses provided comparable fits to the data. In contrast, exploratory analyses suggest a causal impact of both blood-based Aβ biomarkers on NFL, and a reciprocal causal association between t-Tau and NFL.

The absence of clear, empirical support for a causal impact of Aβ on t-Tau, which would be consistent with the amyloid-beta cascade hypothesis, should be interpreted in the context of four considerations. First, our sample was predominately cognitively unimpaired. The proportion of men with mild cognitive impairment (MCI) was 15%.

Causal signals may emerge as the sample ages and the prevalence of MCI increases over time. Second, we relied on plasma biomarkers. While accessible, affordable, and heritable [2], we note that dilution, degradation, and metabolism may introduce variation unrelated to AD-related brain changes. This may limit the predictive validity of these plasma biomarkers to model causation. Ultrasensitive immunoassays and novel mass spectrometry techniques that attempt to address this limitation have begun to show promise in terms of better plasma biomarker measurement [13, 14]. These two limitations are underscored by Coomans et al. [15] who analyzed data from a very small sample of older monozygotic twins with a relatively large number of APOE-ε4 carriers and found significant associations between Aβ-PET and tau-PET. Third, to the extent that plasma Aβ is brain derived, it may nevertheless reflect general health conditions rather than brain amyloid accumulation. Finally, we relied on total tau rather than phosphorylated Tau (p-Tau), which aggregates into neurofibrillary tangles and is therefore likely to be a more relevant indicator of AD pathogenic processes. Indeed, the p-Tau 181, 217 and 231 isoforms have been shown to predict amyloidosis and progression to AD [60]. The genetic variance of these isoforms remains undetermined (including their covariance and direction of causation) with the Aβ and NFL biomarkers. Unlike our results for t-Tau, it is plausible that direction of causation modeling with p-Tau might be consistent with the amyloid cascade hypothesis.

## Conclusion

Notwithstanding the absence of a population-based same-age replication sample, to the extent that that plasma biomarkers are considered informative peripheral indicators of prodromal AD [16-18], our analyses suggest that neither Aβ40 or Aβ42 has any causal impact on t-Tau, when based on community-dwelling sample of men in their late 60s.

## Acknowledgements

The U.S. Department of Veterans Affairs, Department of Defense, National Personnel Records Center, National Archives and Records Administration, Internal Revenue Service, National Opinion Research Center, National Research Council, National Academy of Sciences, Institute for Survey Research, and Temple University provided invaluable assistance in the conduct of the VET Registry. The Cooperative Studies Program of the U.S. Department of Veterans Affairs provided financial support for development and maintenance of the Vietnam Era Twin Registry. We would also like to acknowledge the continued cooperation and participation of the members of the VET Registry and their families

## References

1. Hardy, J.A. and G.A. Higgins, Alzheimer’s disease: the amyloid cascade hypothesis. Science, 1992. 256(5054): p. 184–5.

2. Gillespie, N.A., et al., The heritability of blood-based biomarkers related to risk of Alzheimer’s Disease in a population-based sample of early old-age men. Alzheimer’s & Dementia, 2023. 20(1): p. 356–65.

3. Duffy, D.L. and N.G. Martin, Inferring the direction of causation in cross-sectional twin data: theoretical and empirical considerations [see comments]. Genetic Epidemiology, 1994. 11(6): p. 483–502.

4. Heath, A.C., et al., Testing Hypotheses About Direction of Causation Using Cross-Sectional Family Data. Behavior Genetics, 1993. 23(1): p. 29–50.

5. Neale, M.C., et al., Depression and parental bonding: cause, consequence, or genetic covariance? Genet Epidemiol, 1994. 11(6): p. 503–22.

6. Gillespie, N.A., et al., Direction of causation modeling between cross-sectional measures of parenting and psychological distress in female twins. Behav Genet, 2003. 33(4): p. 383–96.

7. Gillespie, N.A., et al., Modeling the direction of causation between cross-sectional measures of disrupted sleep, anxiety and depression in a sample of male and female Australian twins. J Sleep Res, 2012. 21(6): p. 675–83.

8. Bates, T.C., M. Neale, and H.H. Maes, umx: A library for Structural Equation and Twin Modelling in R. Twin Research and Human Genetics, 2019. 22: p. 27–41.

9. R Development Core Team, R: A language and environment for statistical computing. R Foundation for Statistical Computing, Vienna, Austria. URL https://www.R-project.org/. 2018.

10. Neale, M.C. and L.R. Cardon, Methodology for Genetic Studies of Twins and Families. 1st ed. NATO ASI Series. 1992, Dordrecht: Kluwer Academic Publishers.

11. Boker, S., et al., OpenMx: An Open Source Extended Structural Equation Modeling Framework. Psychometrika, 2011. 76(2): p. 306–317.

12. Akaike, H., Factor analysis and AIC. Psychometrika, 1987. 52: p. 317–332.

13. Blennow, K. and H. Zetterberg, Biomarkers for Alzheimer’s disease: current status and prospects for the future. J Intern Med, 2018. 284(6): p. 643–663.

14. Blennow, K. and H. Zetterberg, Fluid biomarker-based molecular phenotyping of Alzheimer’s disease patients in research and clinical settings. Prog Mol Biol Transl Sci, 2019. 168: p. 3–23.

15. Coomans, E.M., et al., Genetically identical twin-pair difference models support the amyloid cascade hypothesis. Brain, 2023. 146(9): p. 3735–3746.

16. Pérez-Grijalba, V., et al., Plasma Aβ42/40 ratio alone or combined with FDG-PET can accurately predict amyloid-PET positivity: a cross-sectional analysis from the AB255 Study. Alzheimers Res Ther, 2019. 11(1): p. 96.

17. Pérez-Grijalba, V., et al., Plasma Aβ42/40 Ratio Detects Early Stages of Alzheimer’s Disease and Correlates with CSF and Neuroimaging Biomarkers in the AB255 Study. J Prev Alzheimers Dis, 2019. 6(1): p. 34–41.

18. Janelidze, S., et al., Plasma β-amyloid in Alzheimer’s disease and vascular disease. Sci Rep, 2016. 6: p. 26801.

