## Supplement for "Testing the causal impact of amyloidosis on total Tau using a genetically informative sample of adult male twins"

*Biometrical genetical 'twin' modelling to decompose the total variance in each biomarker into additive genetic (A), shared or common (C) environment, and non-shared or unique (E) environmental influences while testing competing causal and non-causal hypotheses.*

We applied the Classical Twin Design (CTD), which relies on twins reared together, to decompose the total variation in each plasma biomarker into additive (A) genetic variance, shared or common environmental (C), and non-shared or unique (E) environmental variance components. This decomposition exploits the expected genetic correlations between MZ and DZ twin pairs; MZ twin pairs are genetically identical, whereas DZ twin pairs share, on average, only half of their genes. Therefore, MZ and DZ twin pair correlations (rA) for additive genetic effects are fixed to 1.0 and 0.5 respectively. The CTD assumes no genotype by environmental interactions or correlations, and random parental mating. It also assumes equal shared environmental effects for MZ and DZ twin pairs, including equality of parental treatment, environmental exposure, and no effects caused by placentation [1]. Given this equal environment assumption, the MZ and DZ twin pair correlations (rC) for shared environmental influences are each fixed to 1.0. All non-shared environmental influences (E), including measurement error, are by definition uncorrelated, so the MZ and DZ twin pair correlation (rE) for these 'E' influences is fixed to zero. This univariate approach can be readily extended to the multivariate case to estimate the size and significance of genetic and environmental influences within and between two or more complex traits, including direction of causation.

Historically, assessing evidence of causality has been challenging without double-blind random case-control experiments or longitudinal designs. In the absence of such data, determining whether an observed association is causal or stems from a correlated liability (where cross-sectional or longitudinal phenotypic associations arise due to correlated, unmeasured background genetic or environmental effects) is difficult. As an alternative to costly genetically informative longitudinal data, we applied an innovative statistical method: direction of causation modeling on cross-sectional, genetically informative data [2, 3]. This approach, which has been applied to various complex behavioral phenotypes [4-6], requires several key assumptions [3]:

1. Members of a twin pair do not have any mutual effect on one another (i.e., no sibling cooperation/rivalry), either within or across variables.
2. The relationship between variables is equivalent for twin 1 and twin 2.
3. Twin pair correlations differ between the variables being studied [7].
4. There are no unmeasured variables influencing both measures, which could inflate correlations arising through the causal influence of one variable on the other.

If these assumptions are satisfied, differences in the patterns of cross-twin cross-plasma biomarker correlations can allow us to falsify strong hypotheses about the direction of causation between two variables measured on a single occasion. The power to do this increases when there are differences in the causes of variation in one biomarker versus another [3].

Figure S1 illustrates this approach. Assuming variable A is best explained by shared (C) and non-shared (E) environmental effects, while variable B is best explained by additive genetic (A), dominant genetic (D), and non-shared (E) environmental effects, we can use Wright's [7] path tracing rules. The 'A-to-B' and 'B-to-A' hypotheses generate different expected monozygotic and dizygotic cross-twin cross-biomarker correlations (e.g., correlation between Twin 1 Biomarker A and Twin 2 Biomarker B), whose goodness of fit can be compared using likelihood-ratio chi-squared tests.

References

1. Scarr, S., *Environmental bias in twin studies.* Eugenics Quarterly, 1968. **15**(1): p. 34-40.

2. Hill, A.B., *The Environment and Disease: Association or Causation?* Proceedings of the Royal Society of Medicine, 1965. **58**: p. 295-300.

3. Heath, A.C., et al., *Testing Hypotheses About Direction of Causation Using Cross-Sectional Family Data.* Behavior Genetics, 1993. **23**(1): p. 29-50.

4. Duffy, D.L. and N.G. Martin, *Inferring the direction of causation in cross-sectional twin data: theoretical and empirical considerations [see comments].* Genetic Epidemiology, 1994. **11**(6): p. 483-502.

5. Neale, M.C., et al., *Depression and parental bonding: cause, consequence, or genetic covariance?* Genet Epidemiol, 1994. **11**(6): p. 503-22.

6. Gillespie, N.A., et al., *Direction of causation modeling between cross-sectional measures of parenting and psychological distress in female twins.* Behav Genet, 2003. **33**(4): p. 383-96.

7. Wright, S., *The method of path coefficients.* Annals of Mathematical Statistics, 1934. **5**: p. 161-215.

8. Martin, N.G., et al., *The power of the classical twin study.* Heredity, 1978. **40**(1): p. 97-116.
