## Supplement Figure S1 for "Testing the causal impact of amyloidosis on total Tau using a genetically informative sample of adult male twins"

**Supplementary Figure S1**. Expected cross-twin cross-trait covariances for monozygotic (MZ) and dizygotic (DZ) twin pairs under the competing uni-directional hypotheses: A causes B; and B causes A.

**Biomarker A causes Biomarker B hypothesis** **Biomarker B causes Biomarker A hypothesis**

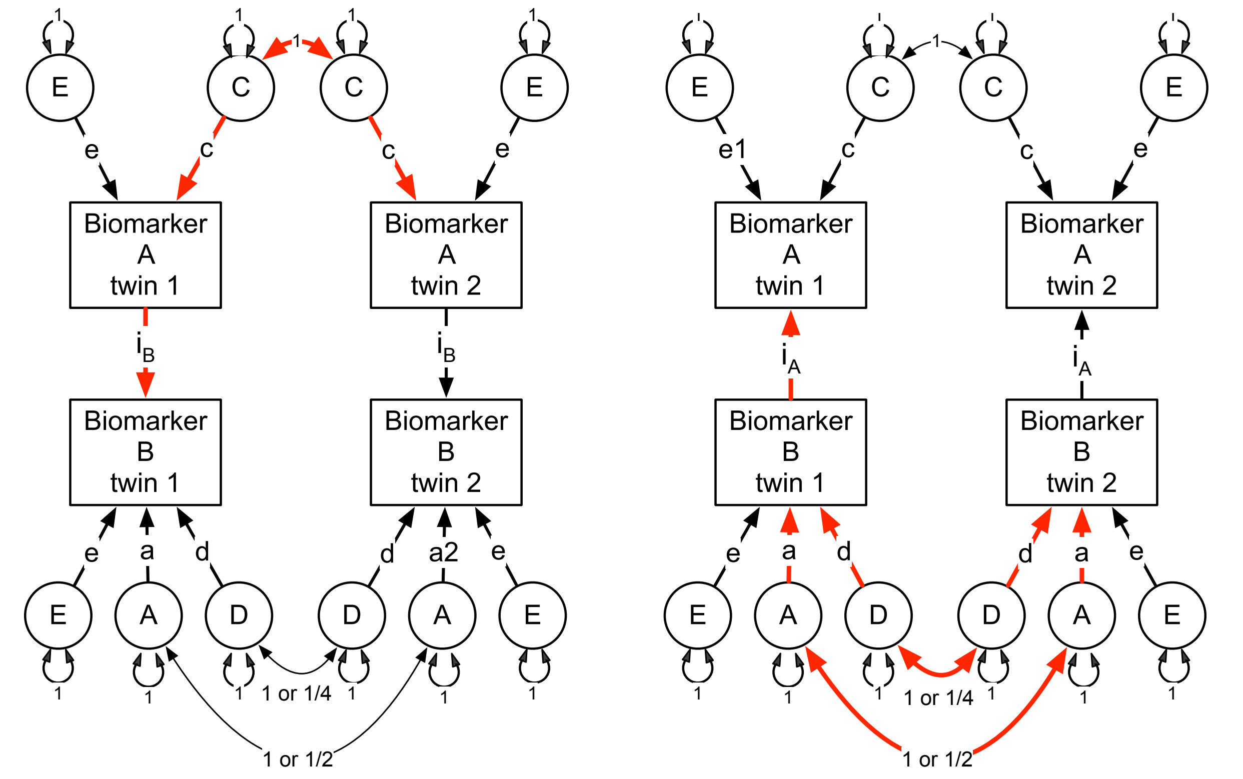


| | |
|---|---|
| Expected MZ and DZ cross-twin cross-trait covariances (red pathway): | Expected MZ and DZ cross-twin cross-trait covariances (red pathway): |
| MZ = $c^2 \times i_B$ | MZ = $(a^2 + d^2) \times i_A$ |
| DZ = $c^2 \times i_B$ | DZ = $(½a^2 + ¼d^2) \times i_A$ |

**Footnote**: A, C, E and D refer to the latent additive genetic, dominant, shared environment and non-shared influences respectively. Double-headed arrows denote latent factor variances and covariances. Means not shown for brevity. Expected cross-twin cross-trait covariances under each uni-directional hypothesis are derived using Wright's (1934) tracing rules, and are bolded in red.

Note that although drawn for illustrative purposes in Supplement Figure S1, we did not model genetic non-additivity or dominance (D). In the CTD, the 'C' and 'D' influences are negatively confounded, and therefore, cannot be modelled simultaneously [8]. Since the sample sizes required to detect 'D' as a source of variation are very large, even for variables measured on a continuous liability scale, we chose to model 'C' influences in all subsequent univariate and multivariate models.
