## Supplement Figure S2 for "Testing the causal impact of amyloidosis on total Tau using a genetically informative sample of adult male twins"

**Supplementary Figure S2**. The non-causal correlated factors model (null hypothesis) and four competing models to account for the associations between both Aβs (Aβ40 & Aβ42) and the t-Tau biomarkers.

(**a**) The null hypothesis - all association via non-causal correlated factors

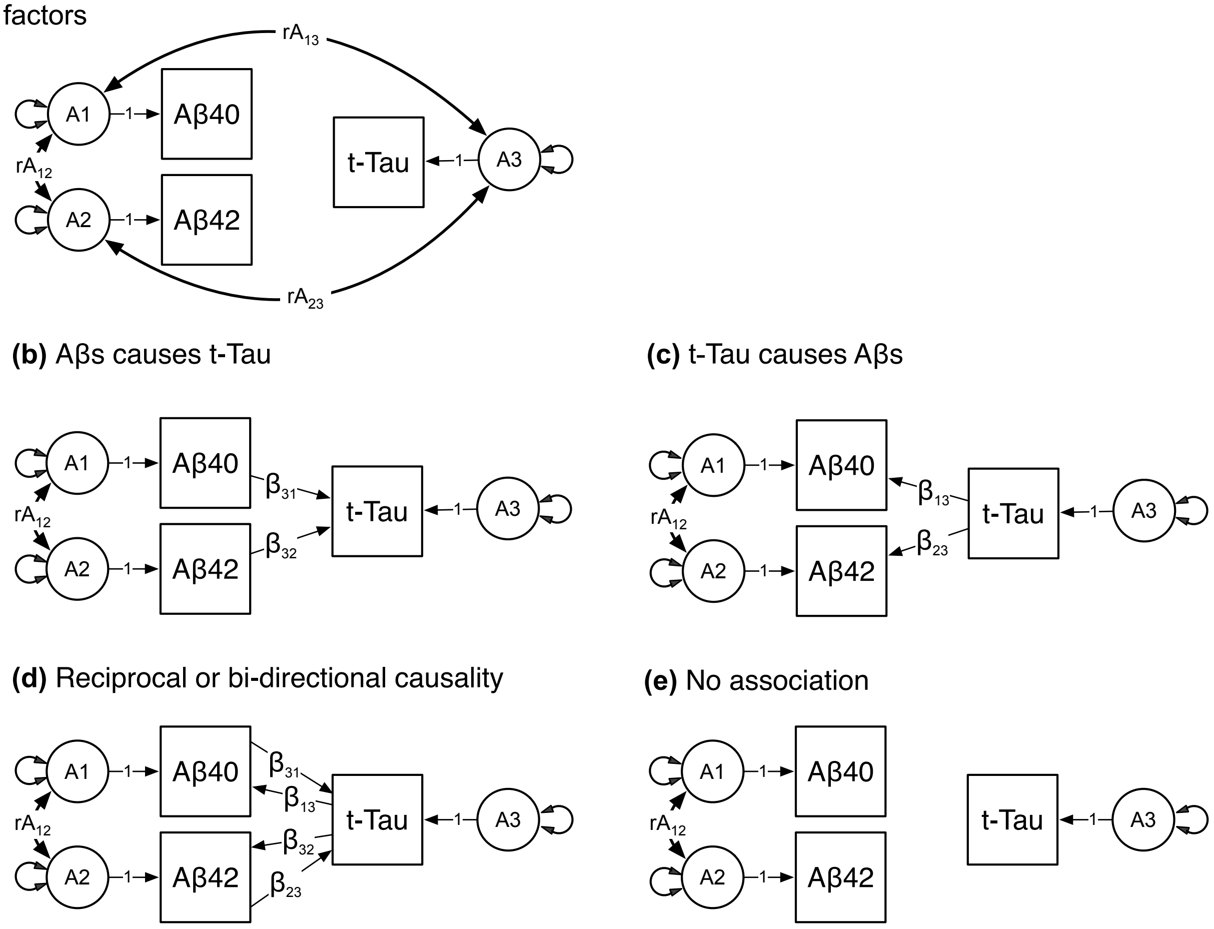


**Footnote**: A1, A2, and A3 refer to the latent additive genetic influences for the Aβ40, Aβ42, and t-Tau biomarkers respectively. Double-headed arrows denote the variances and the non-causal additive genetic covariances. Latent environmental influences, residuals, and means not shown for brevity. Competing models: **(a)** non-causal association stems from correlated additive genetic influences (rA13 & rA23); **(b)** uni-directional hypothesis where Aβs cause t-Tau via the regression coefficients β31 & β32; **(c)** uni-directional hypothesis where t-Tau causes Aβs via the regression coefficients β13 & β23; **(d)** reciprocal or bi-directional causation between the Aβs and t-Tau; and **(e)** no association between the Aβs and t-Tau.
