## Supplement Figure S3 for "Testing the causal impact of amyloidosis on total Tau using a genetically informative sample of adult male twins"

**Supplementary Table S1**. Multivariate model fitting comparisons between the non-causal correlated factors reference model (a), and the two causal (a-b), reciprocal causation (c), and no association (d) models.

| <u>**(i) Both Aβs & NFL**</u> | **ep** | **-2LL** | **df** | **Δ-2LL** | **Δdf** | **p** | **AIC** |
|---|---|---|---|---|---|---|---|
| (a) Correlated / non-causal | 21 | 6420.05 | 3003 | | | | 6462.05 |
| (b) Aβs → NFL | 17 | 6423.97 | 3007 | 3.92 | 4 | 0.4167 | 6457.97 |
| (c) NFL → Aβs | 17 | 6425.97 | 3007 | 5.92 | 4 | 0.2051 | 6459.97 |
| (d) Reciprocal causation | 19 | 6421.17 | 3005 | 1.13 | 2 | 0.5696 | 6459.17 |
| (e) No association | 15 | 6537.34 | 3009 | 117.30 | 6 | 0.0000 | 6567.34 |
| <u>**(ii) NFL & Tau**</u> | | | | | | | |
| (a) Correlated / non-causal | 11 | 4109.36 | 1960 | | | | 4131.36 |
| (b) NFL → Tau | 9 | 4115.62 | 1962 | 6.26 | 2 | 0.0438 | 4133.62 |
| (c) Tau → NFL | 9 | 4113.91 | 1962 | 4.55 | 2 | 0.1029 | 4131.91 |
| (d) Reciprocal causation | 10 | 4110.15 | 1961 | 0.79 | 1 | 0.3736 | 4130.15 |
| (e) No association | 8 | 4141.64 | 1963 | 32.28 | 3 | 0.0000 | 4157.64 |

**Footnote**: Aβs = amyloid-beta 42 & 42, NFL = neurofilament light chain, ep = number of estimated parameters, -2LL = -2 x log-likelihood, Δ-2LL = change in -2 x log-likelihood, Δdf = change in degrees of freedom, AIC = Akaike Information Criteria.
